# MACRO-MOLECULAR CROWDING FAVORS WRITHE IN UNWOUND DNA

**DOI:** 10.64898/2026.04.30.722034

**Authors:** Jin Qian, Zachary Z. Montgomerie, Andrew J. Spakowitz, David Dunlap, Laura Finzi

## Abstract

Genomic DNA is subject to forces and torsion. Some arise mechanically, while others can be entropic, such as those due to crowding within the nuclear environment. Indeed, about 30-40% of the cell is occupied by molecules other than water, and of these, the vast majority are macromolecules. Here, we explore both experimentally and theoretically the interplay between tension, torsion, and macromolecular crowding. Using pharmaceutically relevant crowders of different molecular weights, Dextran 70, and polyethylene glycol (PEG), we observed that macromolecular crowding of unwound, stretched DNA effectively opposed the tension and promoted the formation of plectonemes. A theoretical model representing the equilibrium between *B*- and *L*-form DNA fit to the experimental measurements indicates the contractile tension produced by macromolecular crowding of DNA.

**SIGNIFICANCE STATEMENT:** Distinct DNA conformers are involved in different cellular processes. Genomic DNA is both stretched and unwound by enzymes in a crowded intracellular medium. This can induce conformational changes between extended, twisted and more compact, plectonemic forms. This study explores the effect of macro-molecular crowding on the conformations of DNA subject to tension and torque. Fitting experimental data to a model for the right-to-left-handed DNA transition, we show that macromolecular crowding induces a contractile force that favors DNA writhe and that such force depends both on the concentration and molecular weight of the crowder.

## INTRODUCTION

The negatively charged, double-helical DNA polymer encodes the genetic information of most organisms and propagates through high fidelity, semiconservative replication (1). Alternative structures, or allomorphs, to the familiar right-handed double helix constitute a fundamental level of genomic regulation (2). Besides changes in salt concentrations or pH, mechanical and entropic forces deriving from the activity of proteins and crowding, respectively, can also drive DNA between allomorphs and change the affinity of proteins for specific recognition sites (3-6).

The allomorphs display differences in the pitch of the double helix and the relative sizes of the minor and major grooves that directly impact protein binding and DNA remodeling (7-10). For example, *A*-form DNA has a longer helical rise per base pair and a greater number of base pairs per turn than *B*-DNA (11, 12). Most extraordinary changes are left-handed (*L*-), forms of DNA exhibited by specific sequences in certain conditions (13-15) such as the *Z*-form of poly(dCG) (16, 17). Inversion of helical handedness is much more common than one might expect (18), and increasing evidence links the formation of *L*-DNA in the genome to the negative supercoiling generated during transcription. The extent of *L*-DNA in the genome and the consequences are topics of intense research (14, 19-24), with clear connections to disease (20). Left-handed DNA, which may include melted DNA, forms under less stringent conditions than *Z*-DNA and is much more flexible (persistence length ∼3-4 nm) than *Z*-DNA (persistence length ∼200 nm), or even *B*-DNA (persistence length ∼50 nm) (25). *L*-DNA may be generated (26) in the cell by the activity of DNA-remodeling enzymes, including transcribing RNA polymerase (27) and topoisomerases (28-30). Thus, *L*-DNA may play important roles in a variety of genomic transactions (31), perhaps by forming structures, such as hairpins, that are recognized by specific proteins (32, 33). Indeed, one should imagine that DNA structures and conformations readily change in response to biochemical housekeeping, environment signals, and the transmission of intracellular mechanical stress to the DNA (34-36).

Although the structure of DNA is fundamental in the regulation of genomic function, the configurational dynamics of DNA under tension and torsion are commonly overlooked, and the effect of crowding on transitions between DNA conformers warrants further investigation. To address this fundamental gap, the role of crowding on the conformations of DNA under mechanical tension and torsion was investigated using macromolecular crowders that are relevant in the pharmaceutical industry: Dextran70 (37) and polyethylene glycol (38, 39). Magnetic tweezers were used to twist single DNA molecules (40-42) and record *extension-vs-twist* curves, in various conditions of crowder volume fractions and tension. Crowders shifted the conformational equilibrium of unwound DNA tethers toward plectonemic, writhed forms at the expense of extended, twisted forms, effectively opposing the mechanically applied tension. We compared our data to our recently developed theoretical model (43) that describes the structural transition between right- and left-handed DNA. This model incorporates both the conformational energy of the DNA molecule and a 1D-Ising-like energy function for *B*-to-*L* base pair transitions. The model identified force- and torque-dependent cooperativity that emerges to reproduce the *B*-to-*L* transition, consistent with experimental data from both extension and torque-measurement experiments. Furthermore, this model reproduces the crowding data remarkably well indicating increasing contractile tension at increasing crowder concentrations.

## Materials and Methods

### Preparation of DNA for tethering

DNA tethers for magnetic tweezing experiments were produced by ligation of biotin- and digoxigenin-labeled “handle” fragments to a central fragment. All were PCR amplicons from reactions containing 0.01 ng/µl plasmid template, pDL944 (Figure S1) using 0.33 µM primers (Eurofins, Louisville, KY), 0.025 U/µl Taq DNA polymerase and 0.2 mM nucleotides (New England Biolabs (NEB), Ipswitch, MA). For the central fragment, primers 5’-CCTGGCAGTTCCCTACTC (MF) and 5’-TAACTACGATACGGGAGG (MR) were used to generate a 2707 bp-long DNA fragment with BsaI restriction sites 51 base pairs from the “MF” end and 24 bases from the “MR” end. A 448 bp “handle” fragment for the “MF” end was amplified with primers 5’-CCTTCCCGTTTCGCTC (T1F) and 5’-TCCCGATGGTAGTGTG (T1R) and 10% biotin-11-dUTP (Jena Bioscience GmbH, Jena, Germany) in the dNTP mixture to randomly label “T” positions. Similarly, another 370 bp “handle” fragment for the “MR” end of the central fragment was amplified with primers 5’-AAATCTGGAGCCGGTGAG (T2F) and 5’-AGCAGATTACGCGCAG (T2R) and10% digoxigenin-11-dUTP (ATT Bioquest, Pleasanton, CA) in the dNTP mixture to randomly label “T” positions. All fragments were purified on Uprep microspin columns (Genesee, Morrisville, NC) with Genejet binding and wash buffers (Thermofisher Scientific, Waltham, MA) and eluted in 10 mM Tris-HCl pH 8.0. Then the fragments were digested for 6 hours with 8 units of BsaI-HF (NEB) per µg DNA in the recommended buffer. BsaI is a Type IIS restriction enzyme that recognizes an asymmetric DNA sequence and cleaves beyond it (5’-GGTCTCN’NNNN,). By selecting one PCR primer within 50 bp of a BsaI site and the corresponding, paired primer, over 100 bp beyond that site, the longer cleavage product can be effectively separated from the shorter one by purification on a microspin column (Thermo Fisher Scientific, Waltham, MA). The digests were purified and eluted in Tris-HCl pH 8.0 and mixed at a molar ratio of 2 handle fragments:1 central fragment plus 62,000 units T7 ligase (NEB) per µg of the central fragment for a two-hour ligation. The ligation product was analyzed by gel electrophoresis (Figure S2), purified and eluted in 10 mM Tris-HCl pH 8.0, 1 mM EDTA for immediate use, or storage at -20 °C.

### Preparation of microchambers

Microchambers were assembled with laser-cut, parafilm gaskets between two glass coverslips (44, 45). The volume of a microchamber was about 10□μL. Polyclonal anti-digoxigenin (Millipore-Sigma, St. Louis, MO) was introduced to coat the inner surface of the chamber at a concentration of 8□μg/mL in PBS for 90□min at room temperature. The surface was then passivated with Blocking buffer (PBS with 1% casein, GeneTex, Irvine, CA) for 20□min at room temperature. DNA templates were diluted to 250 pM in PBS and flushed into the passivated microchambers. The solution of DNA templates was incubated in the microchamber for 10□min to anchor the digoxygenin-end of DNA on the anti-digoxigenin coated chamber surface. Then, 20□μL of streptavidin-coated superparamagnetic beads (diluted 1:100 in PBS; MyOneT1 Dynabeads, Invitrogen/Life Technologies, Carlsbad, CA) were flushed into microchambers for attachment to the biotinylated end of surface-tethered DNA templates. After a 5□min incubation, excess superparamagnetic beads were flushed out of the microchamber with 50□μL of PBS.

### Magnetic tweezing experiments

Magnetic tweezing experiments were performed using a custom-built setup described previously (46-48). All measurements were carried out at room temperature. To obtain *extension-vs-twist* curves, intact DNA tethers were first identified by winding 20 turns at low tension and observing the change of their end-to-end distance. Tethers showing a decrease, indicative of plectoneme formation, were selected for further analysis. Then, the DNA tethers were stretched by various amounts by adjusting the distance from the external magnet. The *tension-vs–distance* calibration curve for this setup and bead type has been reported previously (47) and was used to determine the approximate forces applied during the measurements. Finally, to obtain the complete *extension-vs-twist* curve, the extension of the DNA tether was recorded as the magnet was rotated through a range that spanned from -600 to +50 turns at high tension or less at lower tensions to avoid adhesion of the bead to the surface during extensive plectoneme formation.

### Theory for *B*-to-*L* DNA Transition

A full description of the theoretical model can be found in our recently submitted work (43). Here, we provide a summary and highlight key properties. The energetics of the conformation of DNA are captured in the term *βE*_*conf*_ that includes contributions from bending, twisting, and external force and torque. The bending and twisting energies are represented via the twistable worm-like chain (TWLC) model with bend and twist persistence lengths, *l*_*p*_ and *l*_*t*_, respectively (49). However, we define the TWLC energy at the base-pair level; thus, the elastic parameters acquire the superscript *B* or *L* depending on the base-pair state. The force contribution is incorporated as a sum over the work done by the imposed force along the backbone of the molecule, and the torque contribution enters by coupling to the change in linking number, Δ*Lk*, of the DNA molecule.

In addition to *βE*_*conf*_, we incorporate the energetics associated with base-pair structural fluctuations between *B-*DNA and *L-*DNA via a discrete 1D model. This energy, denoted as *βE*_*BL*_, is given by

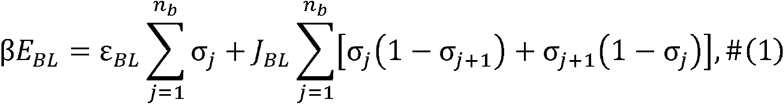

where *n*_*b*_ is the number of base pairs, *ε*_*BL*_ is the energy difference between *B*- and *L*-DNA base pairs and *J*_*BL*_ is the domain-wall energy penalty. We use the structural identifier *σ*_*j*_ to signify the *j*^th^ base pair as *B-*DNA (*σ*_*j*_ = 0) or L-DNA (*σ*_*j*_ = 1). For sufficiently long DNA molecules, the chain ends do not contribute to the overall physical behavior; therefore, we use periodic boundary conditions such that *j* = *n*_*b*_ + 1 = 1.

The statistical behavior of our model is determined from a thermodynamic average (i.e. Boltzman-weighted average) over the molecular conformations, resulting in the partition function. To obtain the partition function, we define a Green’s function that obeys a Schrodinger-like differential equation that can be solved using the methods described by Moroz and Nelson (50). The partition function becomes a sum over all intermediate base-pair orientations and identities and resembles that of a 1D-Ising model, which is solved using the transfer-matrix method.

A surprising prediction of our model is the emergence of a force- and torque-dependent cooperativity Δ*J*(*f,τ*). This additional cooperativity acts to increase the segregation of *B*- and *L*-DNA base pairs into their respective domains and is necessary to explain the value of the torque plateau (–10 pN·nm) that is obtained in torque measurement experiments of the *B-*to-*L* transition (25, 51). The effects and implications of Δ*J* are explored in depth in (43).

Measurements of the persistence lengths for bending, *l*_*p*_^*(B)*^, and twisting, *l*_*t*_^*(B)*^, *B*-DNA produced 53 and 109 nm, respectively, and a base pair rise, 𝓁^*(B)*^, of 0.34 nm (50, 52). The natural twist angle density for *L*-DNA, Ω_*L*_ , and *B*-DNA, Ω_*B*_, are −2π*/*15 and 2π*/*10.5 rad/bp, respectively, therefore, we use ΔΩ_*BL*_ *=* Ω_*L*_ − Ω_*B*_ ≈ −1.0 rad/bp (53). We found that *J*_*BL*_ = 2.3*k*_*B*_*T* at *T* = 300*K* by fitting our model to the experimental data without crowders (43). The value *ε*_*BL*_ = 2.7*k*_*B*_*T* was determined such that the torque plateau corresponded to the experimental value of -10.0 pN·nm (25, 51). Additionally, we found the elastic parameters for *L*-DNA to be *l*_*p*_^*(L)*^ = 7 nm, *l*_*t*_^*(B)*^ = 7 nm, and 𝓁^(*L*)^ = 0.42 nm, which were comparable to previously reported values (25).

## RESULTS

### Macromolecular crowding contracts unwound DNA to favor writhe

Magnetic tweezers were used to wind or unwind single DNA molecules under different levels of stretching tension (54). Under low tension (below ∼0.5 pN), twisting DNA molecules past the buckling point in physiological saline solution produced plectonemes that decreased the end-to-end distance of DNA symmetrically when the molecule was wound or unwound (54-57) (Figure 1A). The slope of either side of the *extension-vs-twist* curve indicates similar average numbers of base pairs in a *B*-form, plectonemic gyre (Figure 1A, purple). However, under slightly higher tension, the extension of unwound DNA does not decrease as rapidly as that of over-wound DNA, and the curve is asymmetric (41, 54, 58) (Figure 1A, orange and yellow). This is due to the coexistence, in unwound DNA, of stretches of plectonemic and extended *L*- and *B*-form DNA. At still higher tension, progressive unwinding produced constant or slightly increasing extension of the DNA tether (59) that persisted until the molecule was completely converted to a left-handed form and further twisting produced plectonemes that rapidly drew the bead to the surface (25, 57, 60, 61) (Figure 1A, blue).

**Figure 1.**
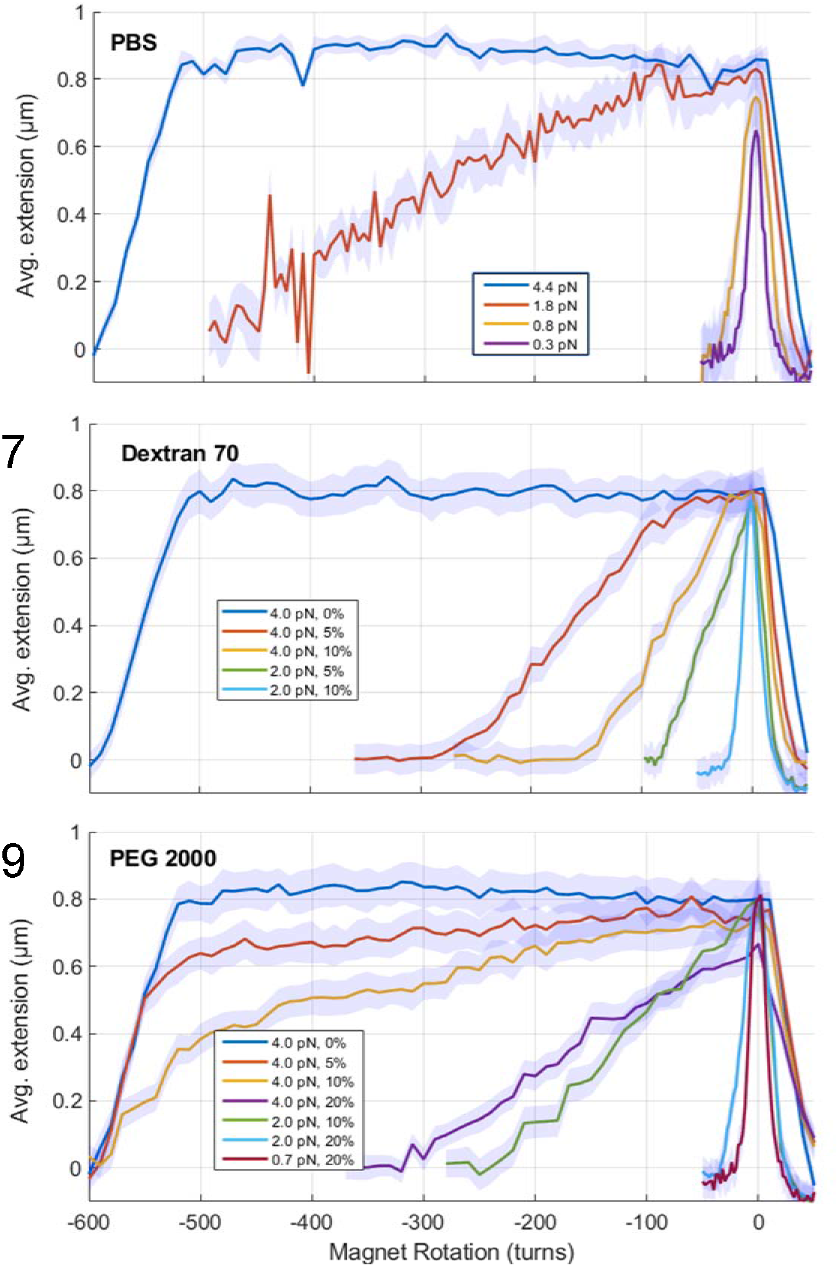
Crowding favored plectonemic, versus extended conformations. Figures show the average tether height (µm) versus the number of turns introduced by the magnetic tweezers. (A) An extension-vs-twist curve in saline buffer. The extension of a DNA tether in the absence of crowder and under different levels of tension was measured as a function of applied twist, turns of the magnet above the microscope stage. Positive twist on DNA tethers under any tension and negative twist on DNA tethers under low tension (purple & yellow) produced plectonemes that reduced the extension of the DNA tether. Under higher tension (red), unwinding caused both twist, that progressively changed the natural right-handed helical DNA tether to a left-handed form, and writhe that reduced the extension of the DNA tether as plectonemes formed. Unwinding the DNA tether under very high tension (blue) progressively converted the molecule to a left-handed helix that finally formed plectonemes beyond -500 turns. (B and C) Both panels include a curve at ∼4 pN without crowder and others with various added percentages of the crowder specified in the legends. The addition of crowder dramatically shifted the DNA conformation toward plectonemes. Similar shifts occurred in molecules under 2 or 0.7 pN of tension.

Macromolecular crowding is known to exert pressure that changes the conformations of macromolecular solutes (62-65). Given the large conformational difference associated with absorbing torsion as writhe or twist, crowding was expected to shift the conformational equilibrium of unwound DNA under tension. To investigate this, magnetic tweezers were used to record *extension-vs-twist* curves in the presence of varying volume fractions of crowders commonly used in the pharmaceutical industry: Dextran 70 (MW=70 kDa), PEG 2000 (MW=2kDa) and PEG 8000 (MW=8kDa) (66). Figure 1B shows a DNA molecule under 4 pN of tension that directly converts to *L*-form upon unwinding in a saline buffer without macromolecular crowder (blue). The extension of the molecule remained constant until more than -500 turns were applied and the molecule was completely converted to *L*-form. Further unwinding produced extensive plectonemes that drew the tethered bead to the surface where the DNA was anchored, similarly to what expected and shown in Figure 1A (blue). Remarkably, adding 5 or 10% Dextran 70 shifted the conformational equilibrium towards plectonemes before -50 turns had been applied and drew the bead to the surface after as little as -300 or -150 turns, respectively (orange and yellow curves). For a DNA tether under only 2 pN of tension, shifting from a solution of 5% to 10% Dextran 70 favored more extensive plectonemes that reduced the end-to-end length of the DNA tether (green and cyan curves). Qualitative comparison of the extension of the DNA tether under a tension of 2 pN in a solution containing 10% Dextran 70, with that of an identical tether in PBS under a tension of 0.8 pN (Figure 1A), indicates that crowding produced a contractile force on the DNA tether of at least (2 – 0.8) pN = 1.2 pN.

A different crowder, PEG 2000, had a similar but less potent effect (Figure 1C). Indeed, a DNA tether under 4 pN of tension in the presence of 5, 10, or 20% PEG 2000 had to be unwound by approximately -550, -500, or only -200 turns to produce plectonemes that reduced the extension to 0.4 µm (orange, yellow and purple curves). The same DNA tether under 2 pN of tension had to be unwound by approximately -120, or just -20, negative turns to reduce the extension to 0.4 µm in the presence of 10 or 20% PEG 2000 (green and cyan curves), respectively. As the tension stretching the DNA was lowered from 2 to 0.7 pN, the *extension-vs-twist* curve in the presence of 20% PEG 2000 returned to a manifestly symmetric (red) curve resembling lower tensions that do not produce *L*-DNA. Under 2 pN externally applied tension, crowding the DNA with 20% PEG 2000 yielded an *extension-vs-twist* curve similar to the one obtained in PBS and 0.8 pN stretching tension (Figure 1A, yellow). Thus, in these conditions, crowding produced a contractile tension in the DNA tether of approximately (2 – 0.8) pN = 1.2 pN. Under 0.7 pN, crowding the DNA with 20% PEG 2000, yielded an *extension-vs-twist* curve perhaps slightly larger than, but similar to the one obtained in PBS and 0.3 pN stretching tension (Figure 1A, purple). By comparison, it appears that 20% crowding produced a contractile tension in the DNA tether of approximately (0.7 – 0.3) pN = 0.4 pN.

Although PEG 8000 made tethered beads more prone to stick to the microchamber and measurements noisier, this crowder too favored writhe over stretched DNA as deduced from the comparison of the curves in the absence (blue and orange) and presence (yellow and purple) of PEG 8000 in Figure S3. The higher molecular weight was more effective than PEG 2000 at contracting DNA. Indeed, Figure S3 shows that adding just 5% PEG 8000 to a DNA tether under 0.7-0.8 pN of tension, produced an *extension-vs-twist* curve similar to one under 0.3 pN in PBS (Figure 1A). Thus, in these conditions, only 5% crowding produced a contractile force on the DNA tether of about (0.75 – 0.3) pN = 0.45 pN (compare Figure S3 yellow and purple and Figure 1A, purple), while 20% PEG 2000 was necessary to produce the same effective decrease in tension in the DNA tether.

These data show that to favor the shift of a DNA tether from extended *L*-to plectonemic form, crowding must oppose the mechanical tension exerted on a DNA tether, reduce the end-to-end extension and create sufficient slack such that mechanically applied torque can be absorbed as writhe instead of twist.

### Theoretical Modeling of Experimental Data

We applied the theoretical model described in (43) and the Material and Methods section to the case of a DNA molecule in the presence of crowding agents. From the experimental results for the DNA extension under both Dextran 70 and PEG 2000/8000, we observed a reduction in the extension as crowder concentration increased. We hypothesized that the normalized extension for increasing crowder concentration could be modeled as an increase in the effective tension within the DNA molecule. To explore this hypothesis, we used a fitting procedure to a model we developed (43) in which the tension was set to the mechanical tension and a set of parameters {P} = {l_p_^(L)^, l_t_^(L)^, 𝓁^(*L*)^, *J*_*BL*_} was fitted. However, in the present study, we used the values of the parameters previously found, and we substituted the actual mechanical tension *f* with the free effective force parameter, *f* ^eff^.

To determine the effective force, we defined and minimized the following cost function

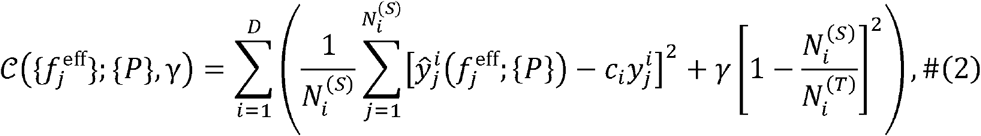

where *c*_*i*_ are fitting coefficients that correct for the uncertainty in the DNA strand’s tether length and are given by

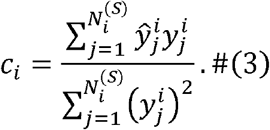

Here, the index *i* spans the number of *extension-vs-twist* curves, D, and the index *j* spans the number of sampled data points for a given *extension-vs-twist* curve, *N*_*i*_^*(S)*^, that were used during fitting. In the present study, D=12 since 7 different conditions of PEG 2000 concentration (%) and 5 conditions of Dextran 70 were used. The first term within the cost function in Eq. 2 is a mean squared loss (MSE) for the error between the model extension *ŷ* = ⟨*z*⟩ / *L*^*B*^ (*L*^*B*^ = *n*_*b*_𝓁^(*B*)^) and the experimental extension *y* at the same value of *σ*_*sh*_. The model extension is parameterized by the set of parameters found in our previous study {*P*}.

The second term is a coverage loss that penalizes the exclusion of experimental data points when N_i_^(S)^ is less than the total number of experimental data points for a given tension N_i_^(T)^. The value of N_i_^(T)^ was determined by requiring both ⟨*z*⟩ */ L*^*B*^ > 0.6 and that the value of the perturbation parameter used to solve our Green’s function is sufficiently small. Data points that did not meet these criteria were excluded (see (43) Supplemental Information for details). We assume the hyperparameter *γ* to be approximately equal to the average squared residual of the MSE. In the present study, we use *γ* = 0.02. Finally, we used the simulated annealing algorithm *scipy*.*optimize*.*dual annealing* (67) to minimize the loss function and optimize fitting coefficients, *c*_*i*_ (Table S1).

Figures 2A and B show the *extension-vs-twist (or - superhelical density)* curves as predicted by our model (lines) and measured using magnetic tweezers (dots) in the presence of crowders Dextran 70 (Figure 2A) or PEG 2000 (Figure 2B). When a molecule was under an applied tension *f* of 4 pN in the presence of Dextran, the effective tension *f*^eff^ fell sharply to 1.6 pN as Dextran concentration was increased from 0% to 5%. This trend continued with *f*^eff^ falling to 1.3 pN as the concentration was increased to 10%. A decrease in the effective force was also seen for a molecule under *f* = 2 pN upon addition of 5 or 10% Dextran. Then, *f*^eff^ was measured to be 1.1 and 0.5 pN respectively. For DNA at *f* = 4 pN in the presence of PEG 2000, the predicted *f*^eff^ was 2.4 pN at 5%. As the PEG 2000 volume percentage was increased to 10% and 20%, *f*^eff^ fell to 1.9 and 1.6 pN, respectively. For DNA stretched by *f* = 2 pN, at 10% and 20% *f*^eff^ became1.5 pN and 0.8 pN, respectively. Finally for the molecule stretched by *f* = 0.7 pN, at 20% *f*^eff^ became 0.5 pN. Thus, increasing the volume fractions of either Dextran or PEG 2000 decreased *f*^eff^ for all applied tensions, though the fall in *f*^eff^ was more gradual in the case of PEG 2000.

**Figure 2.**
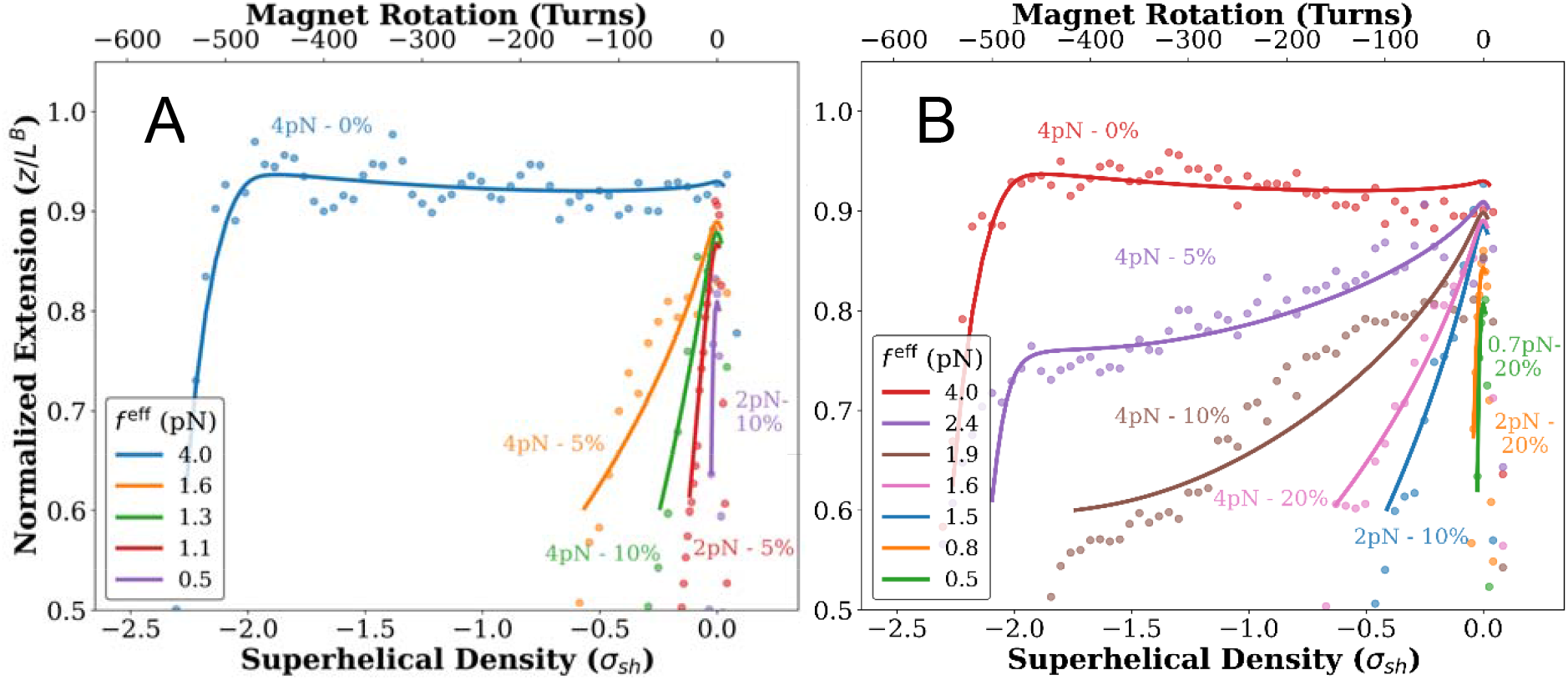
Crowding diminishes the effective tension on DNA molecules under tension and restores plectonemes. Plots show normalized extension versus superhelical density *σ*_sh_. Superimposing experimental data at fixed tensions (dots) and effective tension fit by our model for each experimental condition (solid lines) shows that different concentrations, expressed in %, of crowders drove the DNA tether from extended L-form to plectonemic DNA as it would occur under much lower tensions in the absence of crowder.

The effective tension behavior is summarized in Figure 3 where we plot the difference between the applied force and the effective force (Δ*f* = *f* - *f*^eff^) *vs* crowder concentration for both Dextran 70 and PEG 2000 when the molecule is under applied tensions of 2 pN and 4 pN. From this plot we observe a monotonic increase in Δ*f vs*. concentration for both crowder types and applied tensions.

**Figure 3.**
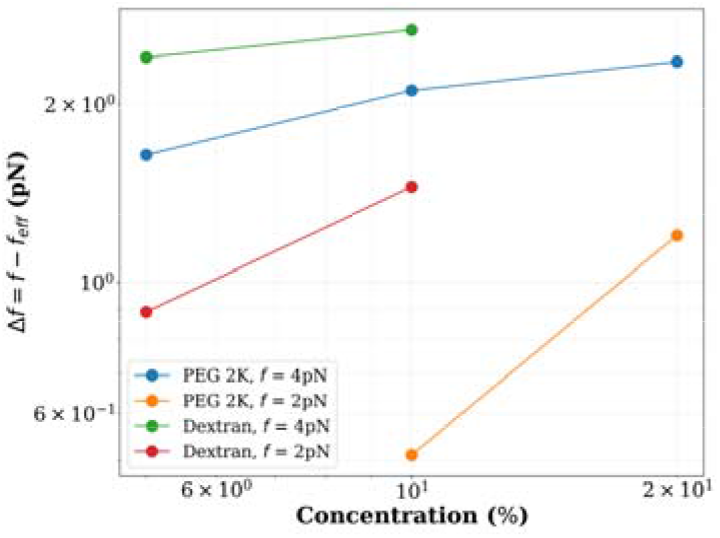
Effective tension difference (Δf = f−feff) vs concentration for solutions of PEG 2000 and Dextran 70, calculated from fitting procedure.

## DISCUSSION

### Macromolecular crowding converts extended *L*-DNA under tension to plectonemes

The torsional state of stretched DNA was manipulated *in vitro* using magnetic tweezers. Under gentle tensions below, or equal to 0.4 pN, the end-to-end extension of single DNA molecules in saline buffer decreased symmetrically whether the molecule was wound or unwound, due to the formation of positive or negative supercoils in B-DNA, respectively. Under slightly higher tensions, between 0.5 and 1 pN, the extension of the DNA decreased more slowly when unwound than when wound due to the force-dependent, partial conversion of *B*-to *L*-DNA. This mixed-phase DNA absorbs torsional stress both as added twist and writhe.

At intermediate tensions between 1 pN to 3 pN, the formation of extended *L*-DNA overtakes that of plectoneme formation. During this *B*-to-*L* transition, the DNA molecule is under a constant torque of -10.0 pN·nm (25, 51), and the extension is an average of the extension of the *B*-DNA and *L*-DNA regions. From our model, we can determine the relative contribution to the writhe from *B*- and *L*-DNA by observing the extension of either form relative to its full length in that form (*i*.*e*. we compare ⟨*z*⟩_*B*_ / *L*^*B*^ to ⟨*z*⟩_*L*_/*L*^*L*^). Additionally, we note that the reduction in extension is a result of both entropic fluctuations and torque-induced writhe. To isolate the effect of writhe and determine the composition of plectonemes in the underwound DNA we compare the relative extensions for *B*- and *L*-DNA both at zero torque and at the torque plateau. Our theory predicts that at zero applied torque and a force of 1.8 pN the relative extension of a pure *B*-DNA molecule is 0.89, while that of a pure *L*-DNA molecule is 0.71. In comparison, we find that at the transition torque at the same force the relative extension of *B*-DNA remains at 0.89, while that for *L*-DNA decreases to 0.60. The significant reduction in the *L*-form extension suggests that during the transition at intermediate tensions, a larger fraction of the writhe is within the *L*-DNA regions.

Under higher tensions, stretched and unwound DNA molecules remained extended absorbing very large numbers of turns as twist before buckling (Figure 1A). However, crowding by Dextran 70, or smaller PEG 2000/PEG 8000 molecules produced transitions in DNA tethers under tension from extended (untwisted) to plectonemic (writhed) conformations (Figures 1B & C). This may be explained conceptually in terms of the excluded volume effect. The volume and entropy available to the crowder (68, 69) in solution are maximized when the (macromolecular) DNA adopts the more compact plectonemic form. The effect was vivid for DNA unwound under 4 pN that readily untwisted in saline buffer but adopted increasingly plectonemic conformations with increasing volume fractions of crowders.

The fact that macromolecular crowders stabilize plectonemic conformations of gently stretched, unwound DNA are supported by the report from Scott *et al*. that DNA unwinding increases with increasing amount of PEG 8000(70), although, their measurement, using Convex Lens-induced Confinement (CLiC) did not produce external tension in the DNA.

Our measurements show that macromolecular crowders effectively exert a contractile force that opposes the externally applied stretching tension on DNA, thus shifting the equilibrium between *L*- and plectonemic DNA towards the latter. In fact, we show that such an effect is a monotonic function of crowder concentration and that the rate at which this shift occurs is higher at lower tensions (2 vs. 4 pN) when the fraction of stretched, *L*-DNA is lower.

### Large macromolecular crowders best stabilized plectonemes

The stabilization of plectonemic DNA was found to be a function of the size of the crowder. Although it is generally believed that relatively smaller crowders better compact DNA (66), PEG 2000, PEG 8000 and Dextran 70 were increasingly effective in reverting extended, unwound DNA to plectonemic DNA conformations in these experiments. This effect is clearly demonstrated by the effective tensions determined from our theoretical model, as shown in Figures 2 and 3. This result might explain observations showing that reduction of the rate of transcription scales with the size of the crowder (71, 72).

### A note on DNA sequence

DNA sequence is likely to tune the effect that crowding has on the conformation of DNA (73), thus altering the affinity of proteins and providing a mechanism for regulating both selectivity and target finding (3-6). The sequence effect on the bending and torsional elasticity of DNA is well illustrated in recent studies of triply H-bonded diamino-purine (DAP)-substituted DNA (55, 57), in which DAP replaces adenine and establishes three H-bonds with thymine throughout the DNA molecule. DAP-DNA adopts an L-form that is considerably more extended than that of the corresponding unmodified sequence (74-76). The effect of macromolecular crowding might be stronger with such DNA for which the conformational shift would portend a greater volume exclusion effect.

### Theoretical Model

This discussion largely ignores denatured bubbles which appear frequently in atomistic simulations of DNA under torsion and tension (77). Whether DNA transits through a denatured state between *B*- and *L*-forms could not be determined in these experiments. Furthermore, simulation of the *B*-to-*L* transition requires accurate force fields and advanced sampling across long timescales (78), which complicates large scale systems, and these single molecule experiments on ∼2600 bp DNA tethers exceed capabilities for atomistic scale simulation. A theoretical framework free of these limitations is therefore needed to understand the interplay between crowding, tension and torsion in the phase transition between *B*- and *L*-DNA. Our recently developed model and its adaptation to the crowded environment address this need and fits the experimental data very well.

### Physiological Relevance

The cell is a crowded environment where 30-40% of the volume is occupied by molecules different from water; of these, about 80% is constituted by macromolecules (66). Given the capability of molecular machinery to apply force and torque (79-81) to DNA and the significant role that left-handed forms of DNA may have in cellular metabolism and disease (14) (note that endogenous Z-DNA sensors like the Zα domain of the ADAR1 protein are likely promiscuous and recognize several left-handed forms, with the possible exception of monoclonal antibodies Z-D11 and Z22 (82)), it is particularly important to characterize how tension and torque affect the *B*-to-*L* transition in the presence of crowders. Mechanisms for mitigating high tensions and torsions that may induce L-DNA formation and interfere with proper physical management of the genome and maintenance of regulatory contacts are, therefore, of great importance (83). Here, we provide experimental and theoretical evidence that macromolecular crowding is one of those mechanisms.

## SUMMARY

In conclusion, our measurements show that macromolecular crowders strongly affect the structure of unwound DNA under tension, favoring plectonemic (writhed) over merely stretched (twisted) DNA. The effect is proportional to the molecular weight of the crowders and is likely to be more pronounced on stiffer sequences, such as those enriched in triply bonded base pairs. The fitting of the experimental data with our theoretical model is remarkable and highlights how such model can be used to understand the interplay between crowding, tension and torsion in the phase transition between *B*- and *L*-DNA.

This study contributes to connect *in vitro* measurements to observations in the crowded cell and to better understand how cellular forces and torques at various locations in the genome and at different times during the cell cycle may flip the handedness of DNA to trigger genomic processes. Future studies will examine the effects of proteins with specific affinity for *L*-DNA.

## AUTHOR CONTRIBUTIONS

J.Q., D.D., Z.Z.M., A.J.S. and L.F. designed research. J.Q. performed single-molecule experiments. Z.Z.M. and A.J.S. developed the theoretical model. D.D., Z.Z.M., A.J.S. and L.F. wrote the manuscript.

## ACKNOWLEDGMENTS

L.F. acknowledges support from grants R01GM084070 and R35GM149296. Z.Z.M. and A.J.S. acknowledge support from the National Science Foundation, Physics of Living Systems Program (PHY-2102726) and from the Alfred P. Sloan Foundation (G-2024-22647).

