## Supplementary material for "MACRO-MOLECULAR CROWDING FAVORS WRITHE IN UNWOUND DNA": SI

### Supplementary Information

#### Table S1

A list of coefficients for fitting one set of model parameters to the ensemble of *extension-vs-turn* data.

| Condition | Crowder | Weight fraction | Tension | Fitting Coefficient |
| --- | --- | --- | --- | --- |
| 1 | PEG 2000 | 10 | 2 | 0.99 |
| 2 | PEG 2000 | 20 | 2 | 0.94 |
| 3 | PEG 2000 | 20 | 0.7 | 0.96 |
| 4 | PEG 2000 | 0 | 4 | 0.89 |
| 5 | PEG 2000 | 5 | 4 | 0.86 |
| 6 | PEG 2000 | 10 | 4 | 0.82 |
| 7 | PEG 2000 | 20 | 4 | 0.88 |
| 8 | Dextran 70 | 5 | 2 | 0.99 |
| 9 | Dextran 70 | 10 | 2 | 0.94 |
| 10 | Dextran 70 | 0 | 4 | 0.89 |
| 11 | Dextran 70 | 5 | 4 | 0.81 |
| 12 | Dextran 70 | 10 | 4 | 0.83 |

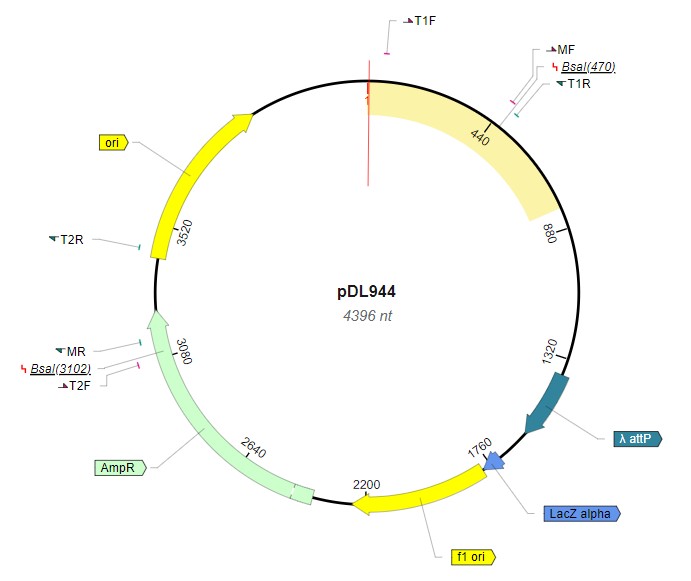

#### Figure S1

Two BsaI restriction sites are present in the pDL944 plasmid. A pair of forward and reverse primers encompassing both sites, MF & MR, were used to amplify a 2707 bp “central” or “main” fragment. Other pairs of forward and reverse primers encompassing only one BsaI restriction site were used to amplify “handle” fragments near MF (T1F & T1R) or MR (T2F & T2R). Restriction of the 2707 bp main amplicon with BsaI clips 24 or 51 base pairs from opposite ends. Restriction of the handle amplicons with BsaI clips 31 or 26 bases from one end of the 370 or 448 bp amplicons respectively. Using microspin columns, these small (< 70 bp) ends can be easily separated from the other 2632, 339, or 422 bp cleavage products. This map was produced using VectorBee (Vector Builder, Chicago, IL).

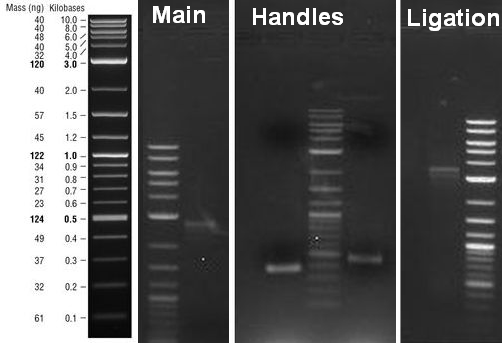

#### Figure S2

Gel electrophoresis was used to verify ligation of the main fragments to the tails. A main fragment of 2707 bp and biotin- or digoxigenin-labeled “handle” fragments of 370, or 448 bp, were produced by PCR (Main and Handles), digested (not shown), and ligated to produce a band near 3.4 kbp corresponding to a biotin- and a digoxigenin-labeled handle joined to opposite ends of the main fragment DNA (Ligation). At left is a reference image of the 1 kbp Plus DNA ladder (New England Biolabs) following electrophoresis.

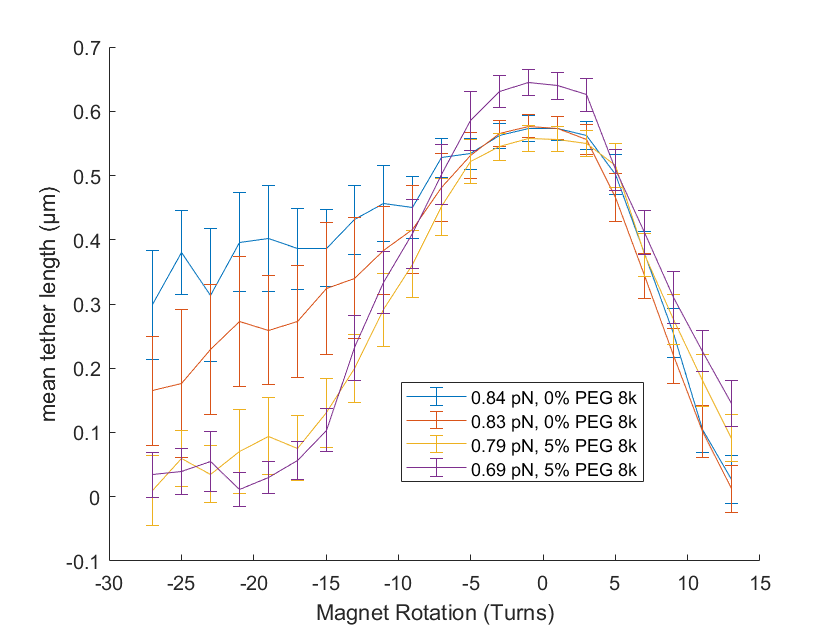

#### Figure S3

Two-tethered beads were stretched in the magnetic tweezer at the indicated tensions as the number of mechanically introduced twists was varied from 15 to –30. Upon winding with no PEG 8000 in the PBS buffer, plectonemes quickly formed and progressively reduced the extension of the tethers. When the molecules were progressively unwound, the negative twist was partially absorbed as twist without forming plectonemes and the tethers remained more extended. When the same molecules were unwound with 5% PEG 8000 in the PBS buffer, the measured tension was slightly less and plectonemes formed to reduce the extension of the tether almost to the same extent observed for positive twisting.
